# Elevated Expression of *srp* RiPPs Across Bacterial Phyla in Marine Sponges

**DOI:** 10.1101/2023.06.09.544420

**Authors:** Samantha C. Waterworth, Evan R. Rees, Chase M. Clark, Skylar Carlson, Ian J. Miller, Melany Puglisi, Jason C. Kwan

## Abstract

We investigated transcriptional activity, at a genome-resolved level, of bacterial communities in two *S. officinalis* and two *I. felix* sponges, both of which are considered high microbial abundance (HMA) sponges that harbor similar bacterial communities. Overlap of metatranscriptional data on genome-resolved metagenomic data showed that genome bins classified within the Chloroflexota and Poribacteria phyla were the most abundant and transcriptionally active. However, abundant bins in general were not the most transcriptionally active, instead less abundant bins of the same phyla were most active indicating that numerical dominance does not imply transcriptional dominance. We found that although some primary metabolic functions appeared upregulated, they were not obviously attributable to any particular bacterial species. However, assessment of transcription levels of biosynthetic gene clusters (BGCs) encoding secondary metabolites revealed a high transcription of ribosomally synthesized and post-translationally modified peptides (RiPPs) in genome bins across diverse bacterial phyla, most of which were likely *srp* RiPPs encoding brominated azol(in)e-containing compounds. However, the ecological role of these compounds remains elusive.

**IMPORTANCE:** Marine sponges and bacteria have formed close associations over several million years with many bacteria performing specialized functions within their sponge host. Previous studies have either assessed the genomes of a given sponge or the expression profile for a sponge holobiont as a whole. Here, we present the first genome-resolved transcriptomic study which gives us a snapshot of the transcriptional activity of individual bacteria in the context of four different sponge holobionts. Using this approach we found that the numerically dominant bacteria were not the most transcriptionally active and that relatively high expression of a ubiquitous biosynthetic gene cluster was evident in many different bacterial phyla in all four sponge samples.

## INTRODUCTION

Over several hundred million years marine sponges (Phylum Porifera) have evolved close associations with microbial symbionts that perform a wide array of functions that can benefit their host or otherwise facilitate a mutualistic or commensal relationship (1, 2). Sponges can be broadly split into two groups: the high-microbial abundance (HMA) sponges and the low-microbial abundance (LMA) sponges. LMA sponges are characterized by the presence of few bacterial cells in their tissues that are often dominated by a single population with Proteobacteria, Bacteroidetes, Planctomycetes, and Firmicutes as indicator phyla (3). The bacterial communities associated with LMA sponges are distinct from the bacterial consortia present in the surrounding seawater and the composition of the associated communities are host specific and distinct between different sponge species (4). Conversely, HMA sponges are characterized by highly diverse communities. Taxonomically distinct HMA sponges have been shown to harbor similar bacterial communities which are often reflective of bacterial species present in the surrounding seawater (4), with bacteria within the Chloroflexi, Acidobacteria, Actinobacteria, PAUC34f, Gemmatimonadetes, BR1093, Poribacteria, AncK6, Nitrospirae, and Spirochaetes phyla as indicator species (3). A recent study investigating the evolution of these two broad classes of sponge holobionts revealed that LMA sponges represent the ancestral form of symbioses, while HMA sponges represent a more modern symbiotic development where host and symbiont have co-evolved in response to the maintenance of more specialized microbial symbionts that play a role in the provision of either primary and secondary metabolites for the sponge host (5).

While there are several studies investigating the functional gene repertoires of sponge holobionts (6–9), there are relatively few studies that have interrogated the transcriptional profiles of these systems (10–12), and none focussing on bacterial symbionts at the genome-resolved level. A functional profile based on differential expression of genes of the HMA *Xestospongia muta* sponge holobiont was generated and showed that the associated bacteria had increased levels of transcripts associated with amino acid biosynthesis, while the host sponge transcribed genes for amino acid catabolism, suggesting a metabolic exchange between host and microbes (13). While this study was not genome-resolved, most transcripts were taxonomically assigned to Firmicutes, Gammaproteobacteria, Cyanobacteria, and Poribacteria, indicating the potential functional importance of these phyla in that particular sponge. Similarly, the tracking of metabolites via NanoSIMS performed in *Plakortis angulospiculatus* (HMA) and the *Halisarca caerulea* (LMA) sponges suggested that associated microbes exchanged primary metabolites with their respective hosts and may be involved in the recycling of host waste products (14). A recent metranscriptomic and metabolomic study of sponge grounds of Arctic-Boreal regions, which are often dominated by HMA sponge species, focussed on sponges of the *Geodia* genus and found that associated bacterial symbionts are responsible for the uptake of dissolved carbon and nitrogen (12). It was further shown that Chloroflexi and Poribacterial symbionts were responsible for the majority of the observed degradation of hydrocarbon compounds and sulfur metabolism (12), as hypothesized previously for these symbionts in Mediterranean sponge *Aplysina aerophoba* (15, 16).

In this study we intended to expand on this knowledge and investigated the transcriptional activity of two HMA sponge holobionts, *Spongia officinalis* and *Ircinia felix*, collected from the Florida Keys. Ultimately, we found no evidence for an outsized role in primary metabolic function in any of the associated bacterial bins, but did identify relatively high transcription of ribosomally synthesized and post-translationally modified peptide (RiPP) biosynthetic genes clusters (BGCs) that are predicted to encode brominated azol(in)e-containing compounds in all four sponges investigated here.

## RESULTS AND DISCUSSION

### Sponge host identity and phylogeny

All sponges collected from the Florida Keys between 2014 and 2015 were initially taxonomically identified through examination of gross morphology. To validate these initial classifications, putative sponge mitogenomes were recovered from available sponge metagenome datasets and annotated for phylogenetic reconstruction using a super-gene tree approach. This study focussed on two putative *Spongia officinalis* sponges (specimens FL20-3 and FL20-9) and two putative *Ircinia felix* sponges (specimens FL2015-9 and FL2015-34) as sufficient quality of metagenomic and metatranscriptomic sequence data were acquired from these sponges. Therefore, we first wanted to confirm the taxonomic identity of these sponges. Phylogenetic analysis of the mitogenome sequences (Fig. 1) showed that the putatively identified *S. officinalis* sponges (specimens FL20-3 and FL20-9) grouped with a reference *S. officinalis* sponge within the *Spongiidae* family alongside *Hippospongia lachne*. The *Irciniidae* and *Verticillitidae* families (also under the *Dictyoceratida* order), aggregated next to *Spongiidae*, with putative *I. felix* sponges from this study (specimens FL2015-9 and FL2015-34) and a reference sequence grouping together into one clade. We additionally included eight other sponges from our group’s collection for taxonomic validation: FL2015-44 was confirmed to be *N. proxima* within the order *Haplosclerida*, as this grouped with other *Haplosclerida* sponges–*Amphimedon compressa*, *Callyspongia plicifera* and *Xestospongia muta*. FL2015-37 was confirmed to likely be *Spheciospongia vesparium* as this grouped with other members of the *Clionaidae* family (*Cliona varians*) along with putative *S. vesparium* sponges, FL2015-4 and FL2015-5.

**Figure 1.**
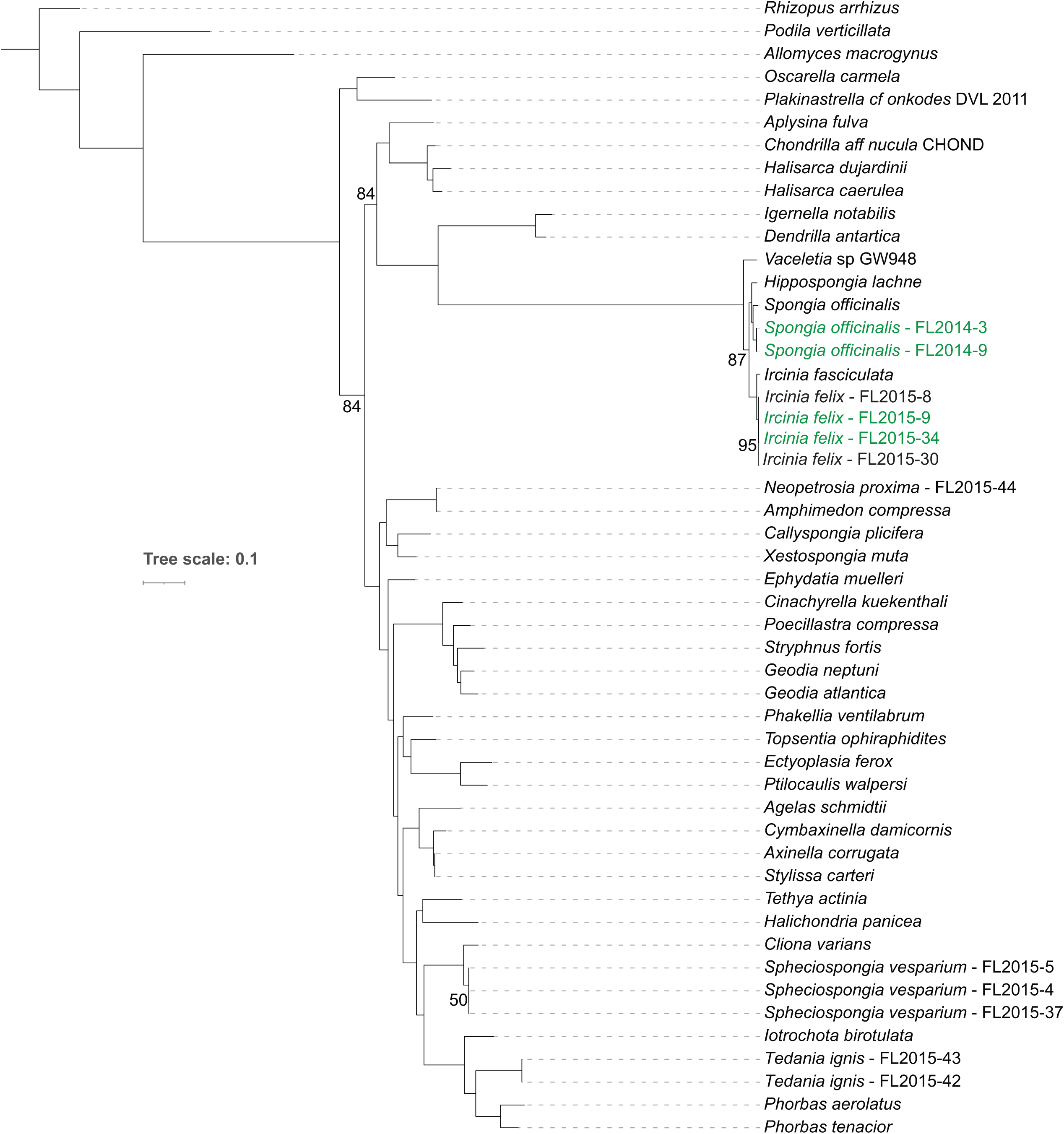
Phylogeny of sponges in this study based upon a Bayesian inference tree using a partitioned data model of nine protein-coding genes from the sponge mitogenomes. Determined posterior-probabilities are 100 unless otherwise noted. Sponge specimens from this study are highlighted in green.

### Bacterial community structure

Bacterial community structure was preliminarily assessed using 16S rRNA gene amplicon sequence analysis with Operational Taxonomic Units (OTUs) clustered at a distance of 0.03. The majority of the top 10 most numerically dominant OTUs (Fig. 2) were matched to bacterial taxa known to be associated with marine sponges. (Table 1). ANOSIM analysis comparing the communities associated with *S. officinalis* and *I. felix* sponges showed the bacterial communities were significantly different (*p* = 0.01) but a low R-statistic (R = 0.154) indicated that differences in the bacterial communities are not likely a result of the sponge host identity. Two sponges from each sponge species (4 samples in total) were chosen for further shotgun metagenomic analysis (indicated by asterisks in Fig. 2) on the basis of DNA yield and the fact that their microbiomes were not overly dominated by *Synecococcus*, which might risk domination of resulting read depth.

**Figure 2.**
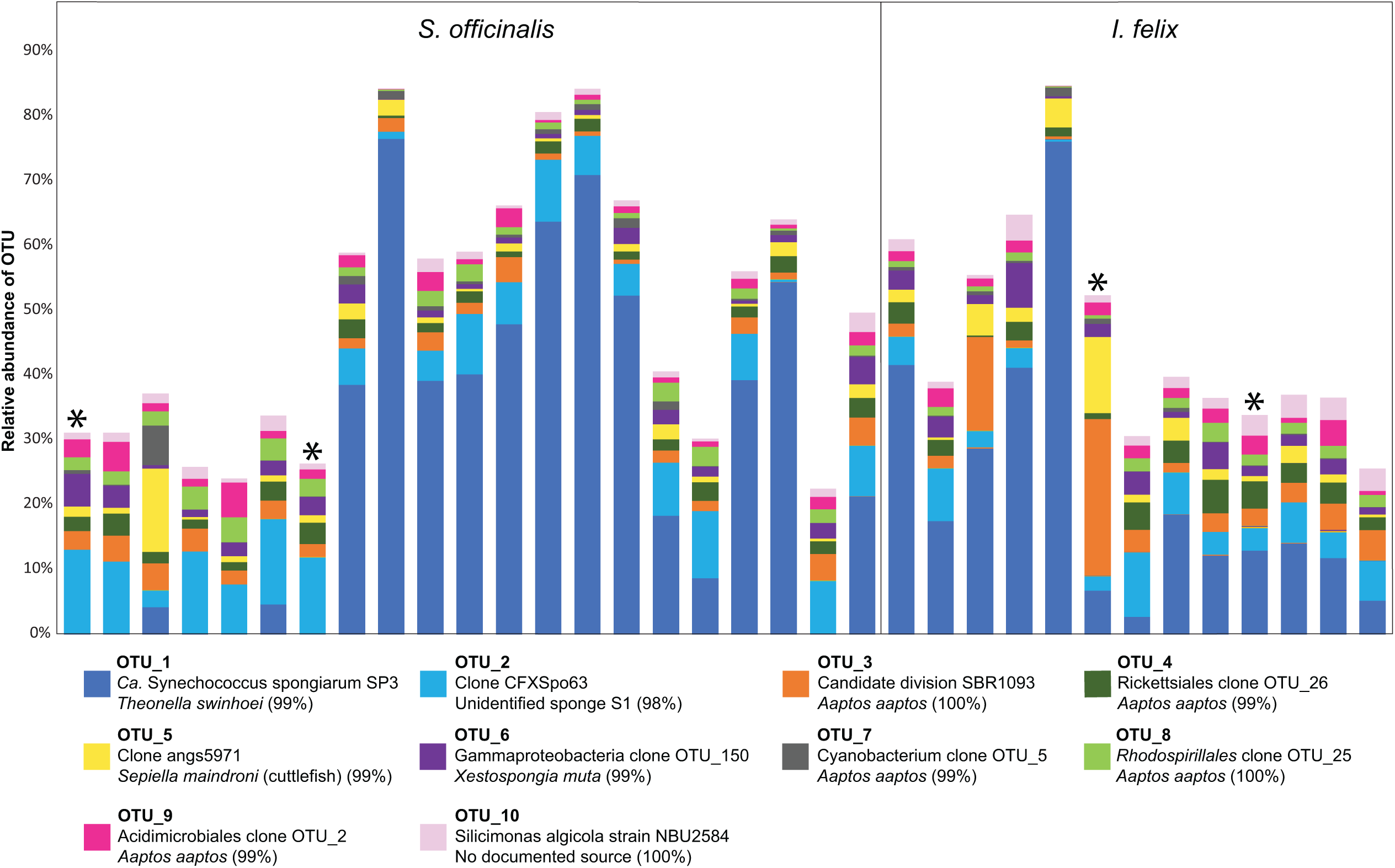
The relative abundance of the top ten most abundant OTUs in *S. officinalis* and *I. felix* sponges. A colored key is provided for OTU identification, where OTU, closest match, isolation organisms, and percent identity are provided. Asterisks indicate samples that were used for metagenomic analyses.

**Table 1.** Closest matches of top 10 most abundant OTUs in *S. officinalis* and *I. felix* against the NCBI nr database

| OTU | Organism | % identity | % cov | Evalue | Source | Accession |
| --- | --- | --- | --- | --- | --- | --- |
| OTU1 | Candidatus<br>Synechococcus<br>spongiarum SP3<br>clone SP3-8 | 99.05 | 100 | 2e-43 | *Theonella swinhoei*<br>marine sponge | KP792324.1 |
| OTU2 | Uncultured<br>bacterium clone<br>CFXSp63 | 99.04 | 100 | 2e-43 | Unidentified marine<br>sponge S1 | GU971200.1 |
| OTU3 | Uncultured<br>candidate division<br>SBR1093 bacterium<br>clone OTU_23 | 100 | 97 | 2e-43 | *Aaptos aaptos* marine<br>sponge | MT464550.1 |
| OTU4 | Uncultured<br>Rickettsiales<br>bacterium clone<br>OTU_26 | 100 | 100 | 2e-42 | *Xestospongia muta*<br>marine sponge | MT464780.1 |
| OTU5 | Uncultured<br>bacterium clone<br>angs5971 | 99.02 | 98 | 3e-42 | *Sepiella maindroni*<br>(cuttlefish) | KR303237.1 |
| OTU6 | Uncultured<br>Gammaproteobacte<br>ria bacterium clone<br>OTU_150 | 98.98 | 97 | 2e-39 | *Xestospongia muta*<br>marine sponge | MT464740.1 |
| OTU7 | Uncultured<br>cyanobacterium<br>clone OTU_5 | 99.04 | 100 | 2e-43 | *Aaptos aaptos* marine<br>sponge | MT464617.1 |
| OTU8 | Uncultured<br>Rhodospirillales<br>bacterium clone<br>OTU_25 | 100 | 100 | 6e-44 | *Aaptos aaptos* marine<br>sponge | MT464566.1 |
| OTU9 | Uncultured<br>Acidimicrobiales<br>bacterium clone<br>OTU_2 | 98.08 | 100 | 1e-41 | *Aaptos aaptos* marine<br>sponge | MT464529.1 |
| OTU10 | *Silicimonas algicola*<br>strain NBU2584 | 100 | 100 | 6e-44 | Unknown | MT636642.1 |

### Metagenomic analyses of *S. officinalis* and *I. felix* sponge bacterial communities

Two metagenomic sequence datasets were generated from each of the two different sponge species (*S. officinalis* and *I. felix*) for a total of four metagenomic datasets. Clustering of contigs into kingdoms showed that bacterial contigs comprised 70–80% of all contigs in *S. officinalis* and *I. felix* sponges (Fig. 3). All bacterial contigs were then further clustered into putative genome bins. We recovered 81 high quality, 83 medium quality, and 109 low-quality bins from the *S. officinalis* sponges and 32 high quality, 100 medium quality, and 144 low-quality bins, from the *I. felix* sponges (Table S1), according to MIMAG standards (17).

**Figure 3:**
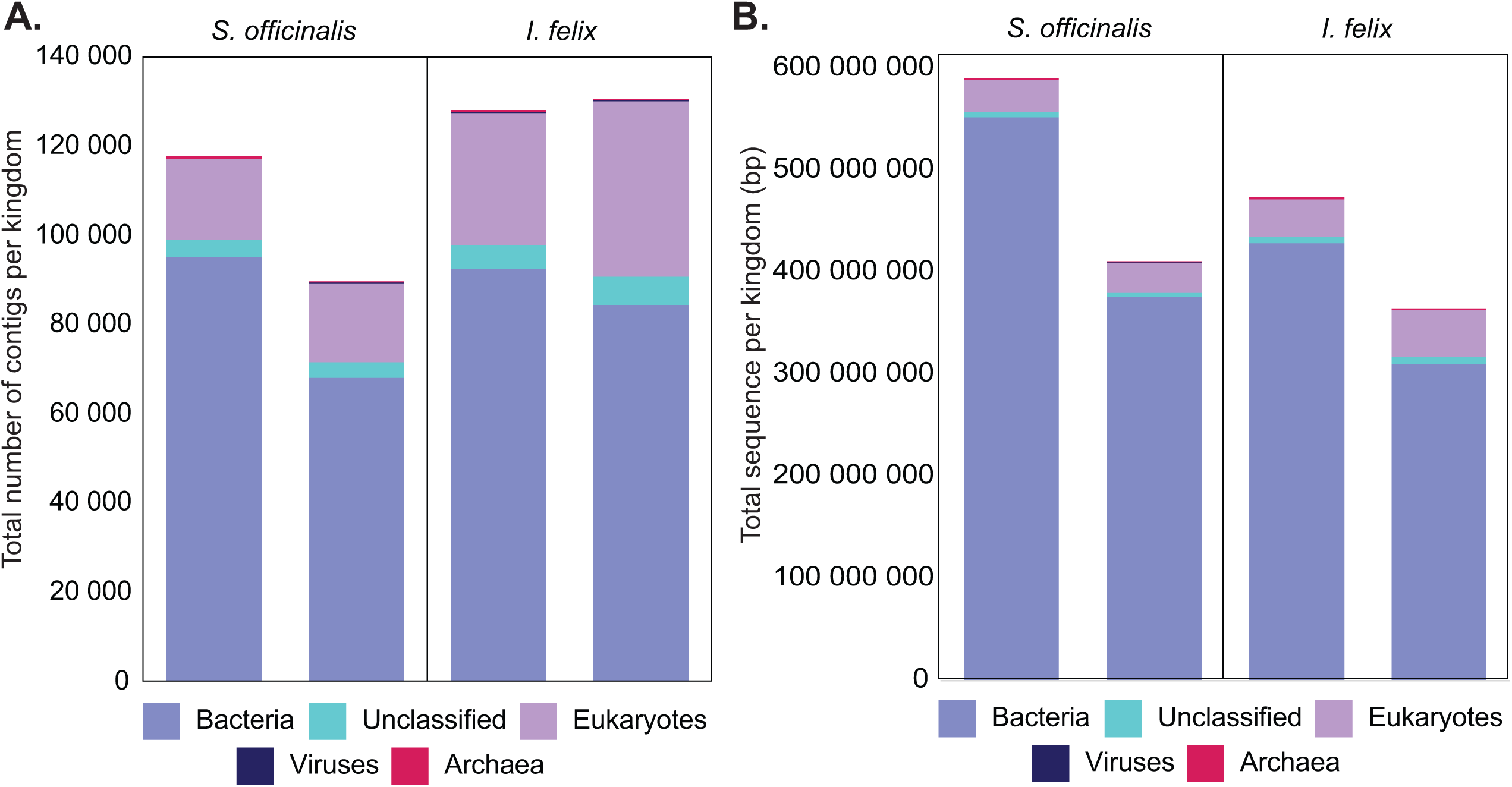
Distribution of **A.** number of contigs and **B.** total length of contigs per kingdom.

Genomic bin abundance was estimated using coverage as a proxy. It was revealed that there was a high abundance of bins classified as Proteobacteria and Chloroflexota in *S. officinalis* sponges, and a high abundance of Poribacteria and Chloroflexota bins in the *I. felix* sponges (Fig. 4A). Considering only the high and medium quality bins, the most abundant bacteria were Poribacteria in the *I. felix* sponges and Chloroflexota in *S. officinalis* sponges (Fig. 4B). Poribacteria were not detected in the 16S amplicon analysis (Fig. 2). However, this inconsistency between the 16S amplicon analysis and metagenomic study is not surprising, as Poribacteria are often missed in these amplicon analyses due to a mismatch between traditional amplicon primers and the Poribacteria 16S sequences (18–20).

**Figure 4.**
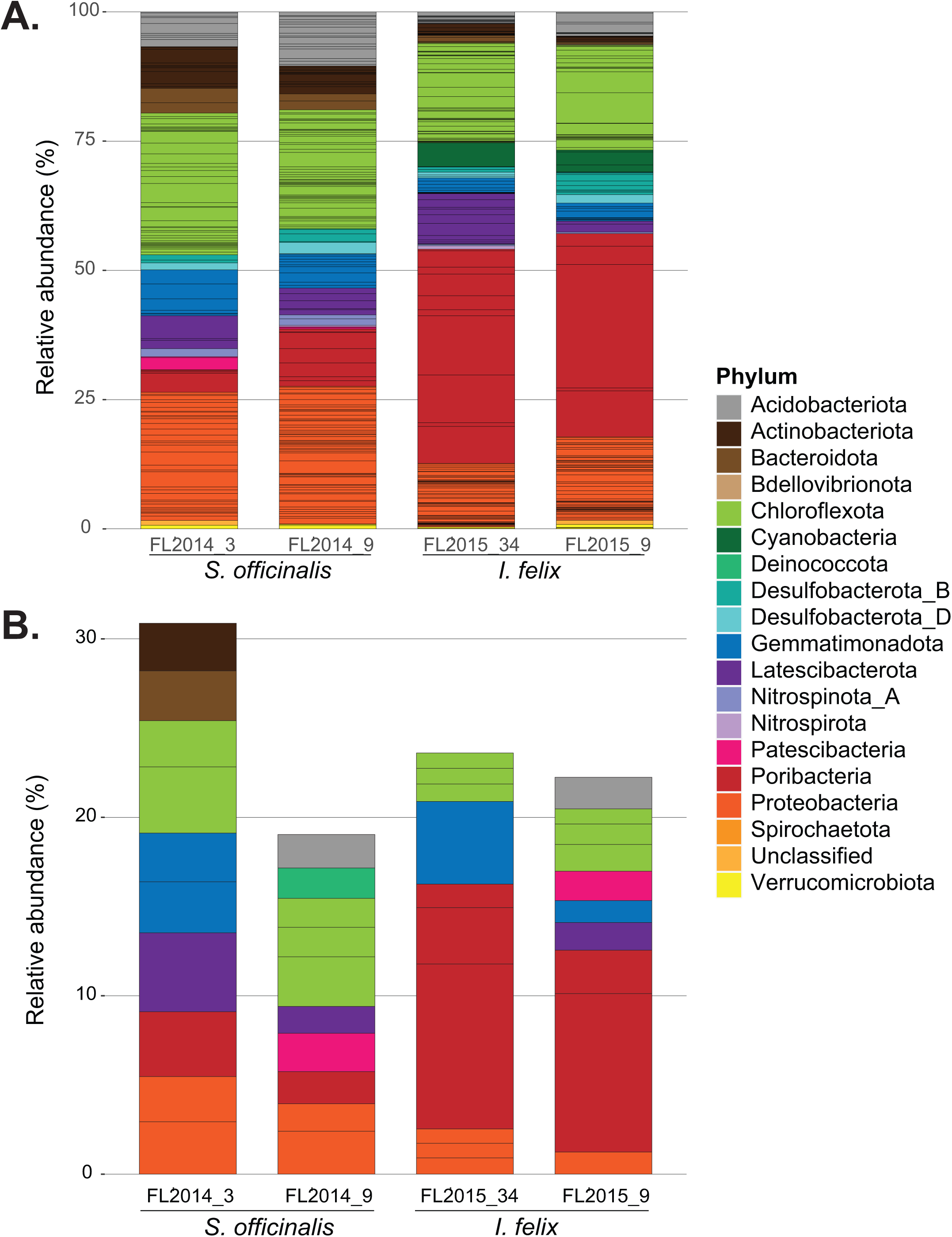
Taxonomic classification of bins per sponge sample scaled by relative abundance in the respective samples. **A.** All bins clustered in *S. officinalis* and *I. felix* sponges and **B.** The top ten most abundant high/medium quality genome bins in each sponge sample. Each block within the bar graph represents a genome. The height of each block represents the relative abundance of that genome bin within the respective samples. Bins were taxonomically classified using GTDB-Tk and colored according to phylum.

### Metatranscriptomic analyses of *S. officinalis* and *I. felix* sponge bacterial communities

We then aimed to investigate the transcriptional activity of the sponge bacterial communities in *I. felix* and *S. officinalis*. Expression of individual genes was calculated as the total expression of all genes in a given sample, and the average expression per bin was calculated as an average of relative expression of all genes within that bin. Assessment of expression levels per bacterial cell (i.e expression corrected for genome abundance) revealed that the more abundant genome bins were not the most transcriptionally active. However, both the most abundant and the most transcriptionally active bacterial genome bins were classified within the Poribacteria, Chloroflexota and Proteobacteria phyla (Fig. 5, Table S1).

**Figure 5.**
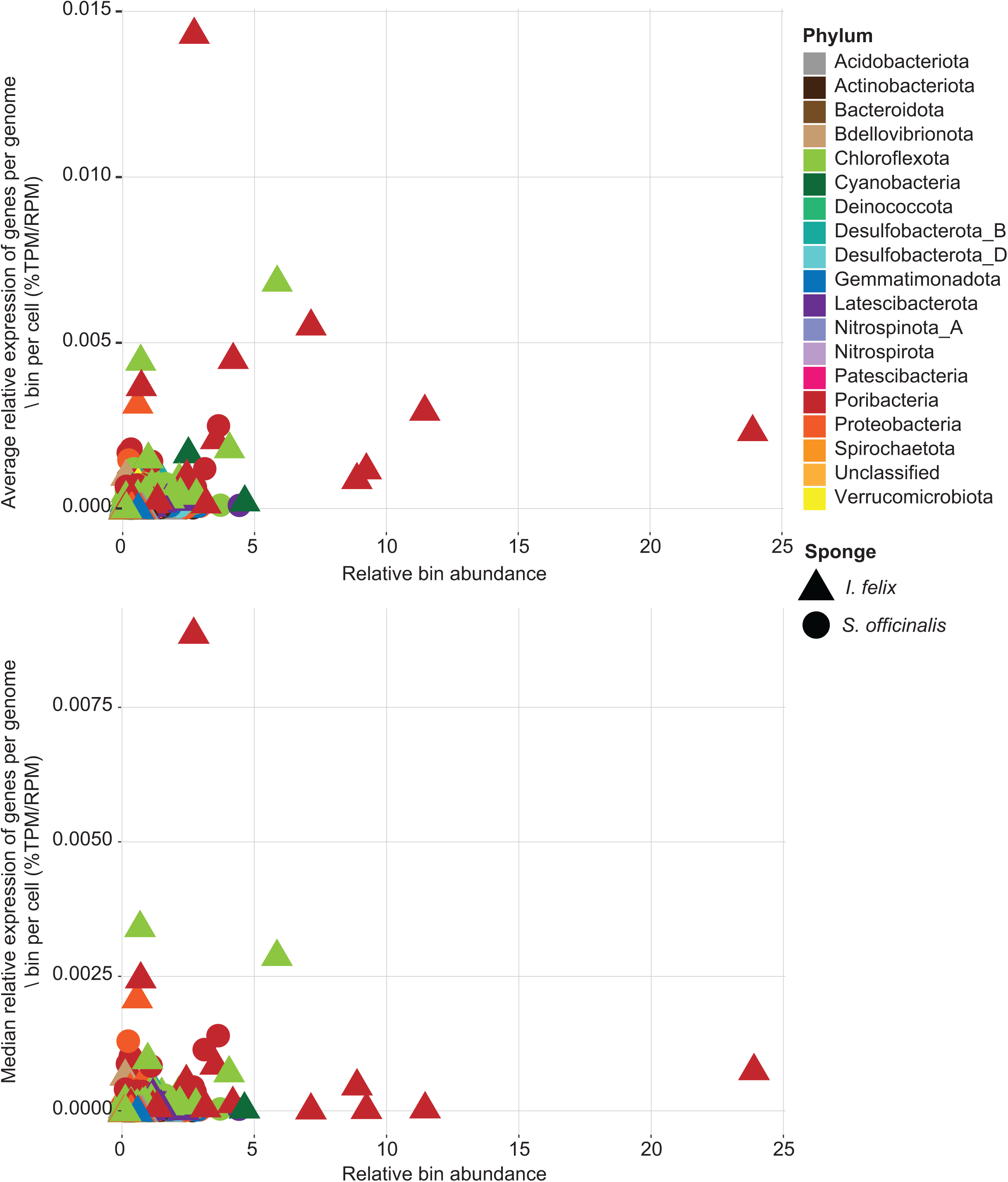
Genome bin abundance relative to transcriptional activity **A.** average relative gene expression levels per bin per bacterial cell and **B.** median relative gene expression levels per bin per bacterial cell. Taxonomic classification of genome bins at the phylum level is indicated by a colored key. The sponge from which the bin was extracted is indicated by shape.

### Primary metabolic function of sponge-associated bacteria

All genes (binned and unbinned) were initially annotated against the KEGG database. Inspection of median relative gene expression per KEGG pathway revealed that the most highly transcribed genes are associated with the ribosome, RNA polymerase, bacterial infection, and valine, leucine and isoleucine biosynthesis (Table S2). This may be suggestive that associated bacteria may help produce leucine and isoleucine for the sponge host, as they are essential amino acids for Porifera (21). However, very few genes were successfully annotated against the KEGG database, with an average of only 33% of genes annotated per sample (Table S3). All genes were additionally annotated against the COG database, where an average of 62% of all genes were successfully annotated. Considering the median expression per COG category per sponge sample, we found that genes associated with Extracellular structures (Category W), Translation, ribosomal structure and biogenesis (Category J) and Energy production and conversion (Category C) were expressed at relatively high levels (Table S4). Inspection of median and average expression per COG category in individual MAGs revealed no correlation with any taxonomic level suggesting that no single taxon appeared to have an enrichment in any given COG category suggesting that no bacteria have a specialized or outsized primary metabolic roles in the sponge holobiont (Fig. S1).

### Secondary metabolic function of sponge-associated bacteria

Biosynthetic gene clusters (BGCs) were identified in bacterial contigs using antiSMASH. A total of 6044 biosynthetic gene clusters (BGCs) were recovered from the four samples (Table 2).

**Table 2:** Number of BGCs recovered from each of the four specimens categorized by BGC type

|  | *Spongia officinalis* |  | *Ircinia felix* |  |  |
| --- | --- | --- | --- | --- | --- |
| Type | FL2014_3 | FL2014_9 | FL2015_9 | FL2015_34 | Total |
| Other | 986 | 1342 | 614 | 1041 | 3956 |
| **Terpene** | 183 | 259 | 148 | 207 | 797 |
| **PKSI** | 134 | 196 | 109 | 127 | 566 |
| **RiPP** | 110 | 171 | 114 | 141 | 536 |
| **NRPS** | 39 | 53 | 23 | 39 | 154 |
| **PKS_Other** | 9 | 9 | 4 | 8 | 30 |
| **Saccharide** | 1 | 2 | 1 | 1 | 5 |
| **Total** | 1462 | 2032 | 1013 | 1537 | 6044 |

The most abundant BGC types were those classified as “Other”. BGCs classified as terpenes, Type I PKS, and RiPPs were similarly abundant in all four specimens.“RiPP” BGCs had the greatest average relative expression in all four sponges, with notable expression levels of BGCs classified within the of “NRPS”, “PKS_Other” and “Other” BGC types in *I. felix* FL2015_9 (Fig. 6).

**Figure 6.**
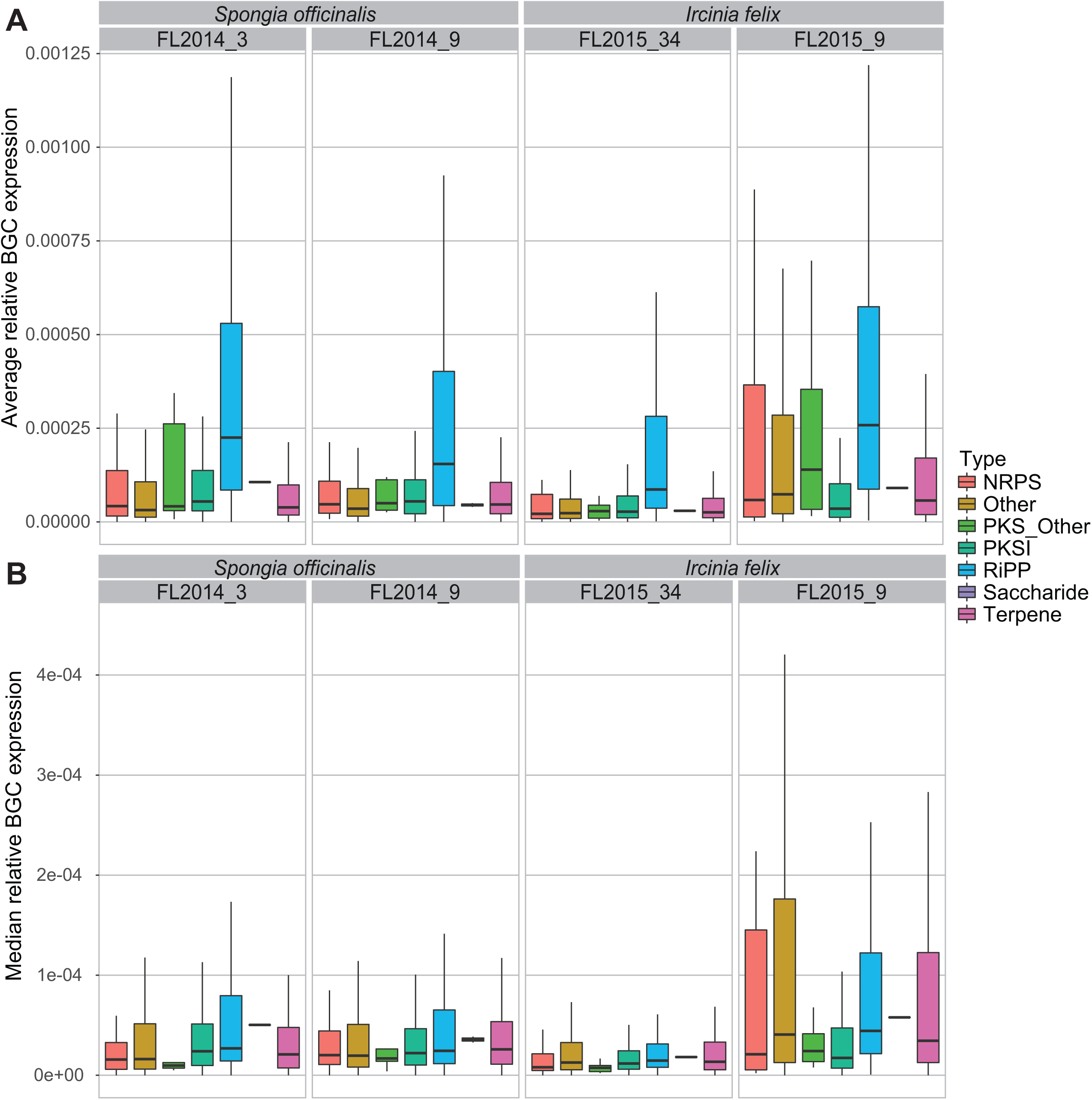
The average (**A**) and median (**B**) relative expression per biosynthetic gene cluster (BGC) pathway type in each of the four sponge samples. It was noted that RiPP BGCs had higher average relative expression in all samples.

Next, we assessed similarities between the biosynthetic gene clusters using BiG-SCAPE (22). Visualization of the resultant networks (Fig. S2), with expression data included, revealed there were six notable networks. Two networks included BGCs that were comparatively highly transcribed relative to other identified BGCs. The first network consisted of 109 BGCs classified as “RiPPs” and the second consisted of 12 BGCs classified as “Other” (Fig. 7A). Additionally, there were four smaller networks where certain BGCs appeared to be highly transcribed, one of which included BGCs classified as “RiPPs” and three classified as “Other (Fig. 7B).

**Figure 7.**
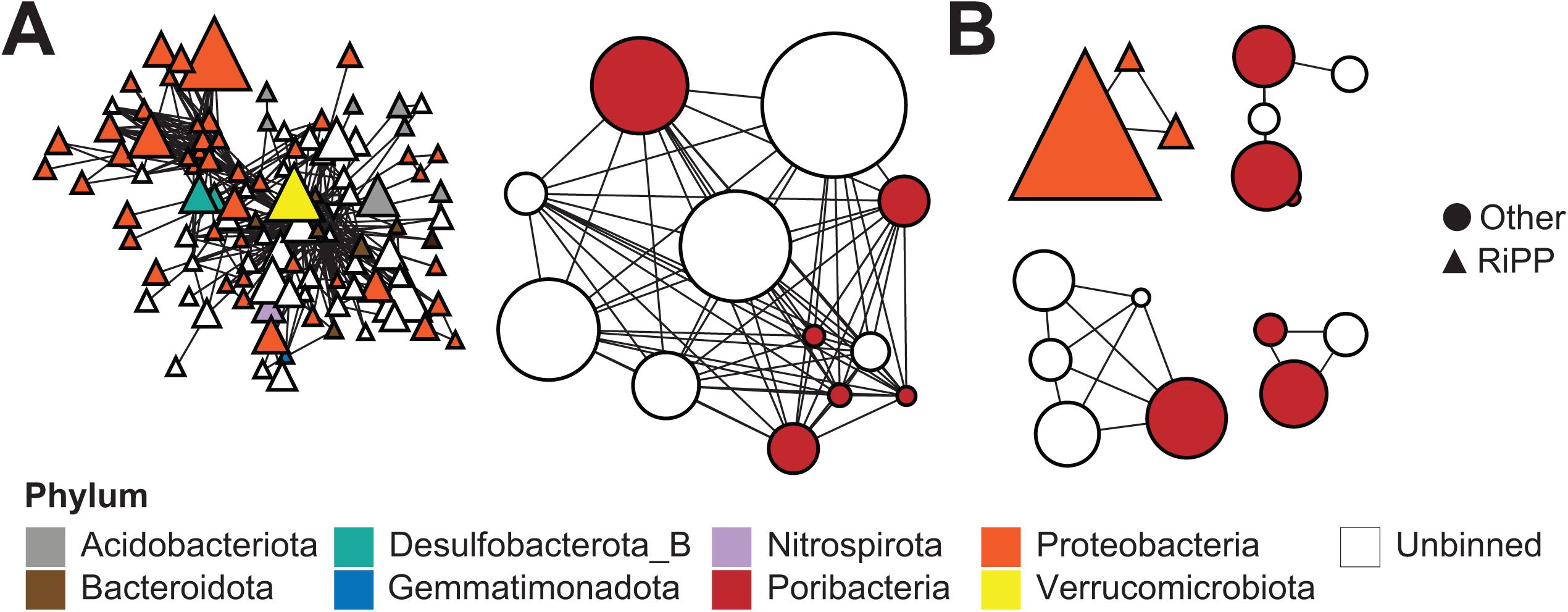
Biosynthetic gene cluster networks were selected for further investigation as they included many related BGCs (**A**) or BGCs included in the network were highly transcribed (**B**). Color of nodes indicates the taxonomic classification of the genome bin in which the BGC was found. The shape of the node indicates the classification of the BGC type. Size of network nodes is indicative of average relative expression of the BGC.

First, we investigated the four “Other” BGC clusters that were highly expressed. These BGCs were present either in bins classified as Poribacteria or on contigs that remained unbinned. Each of the networks were visualized using clinker (23) (Fig. S3) and a representative cluster was chosen for illustrative purposes (Fig. 8).

**Figure 8.**
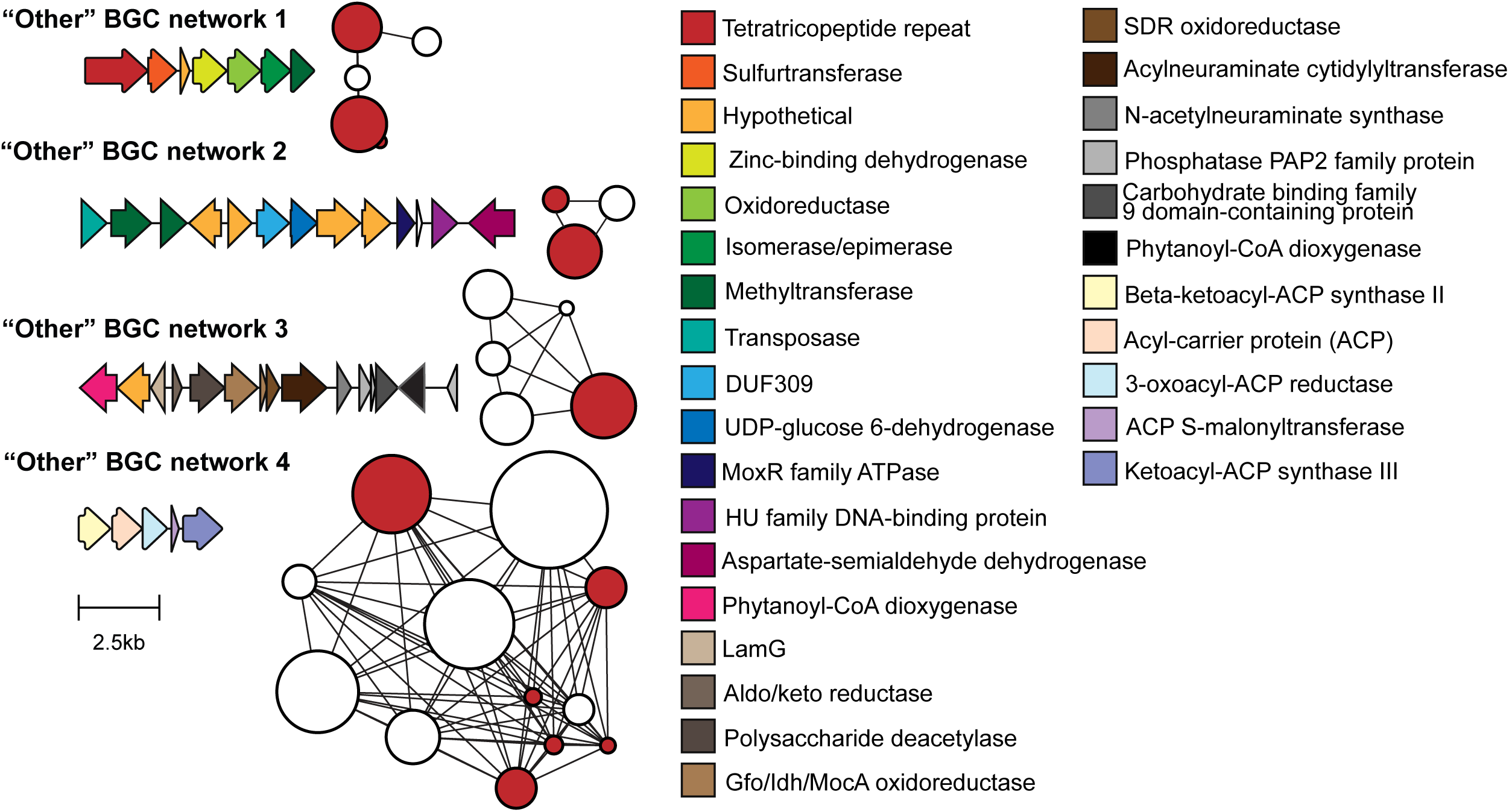
Representative BGCs from each of the four most highly transcribed networks of BGCs classified as “Other”. A color key is provided for gene annotations and a scale bar is provided for approximation of gene size. Size of network nodes is indicative of average relative expression of the BGC.

Homologous gene clusters were searched for in 141 reference Poribacterial genomes using cblaster (24). The BGCs from networks 1, 3, and 4 were found to be highly conserved and present in 16 (11%), 3 (2%) and 45 (32%) of the reference Poribacterial genomes respectively. BGC network 2 did not appear to have any close homologs in the 141 reference genomes tested. It is unclear what compound(s) the BGCs in Networks 1, 2, and 3 may produce. Inspection of the similar clusters identified in the 45 reference genomes suggested that Network 4 BGCs are likely involved in fatty acid biosynthesis rather than the production of any secondary metabolite, as closest matches correlate with the *fab* gene operon (i.e. *fabH* - ketoacyl-ACP synthase III, *fabD* - ACP-S-malonyltransferase, *fabG* - 3-oxoacyl-ACP reductase, *fabF* - beta-ketoacyl-ACP synthase II, acpP - acyl-carrier protein).

Next, due to the high average transcription levels across the four samples (Fig. 6) and the large number of BGCs that shared at least 30% similarity (Fig. 7), we began a deeper investigation into the shared RiPP BGCs. A total of 536 RiPP BGCs were identified across the four sponges (Table 2). The majority of RiPPS were identified in bins classified within the Proteobacteria phylum (238, 44%) and on unbinned contigs (193, 36%) (Table S5). Closer inspection of the 536 BGCs at a gene-resolved level (Table S6) revealed that in most cases elevated average relative expression levels were caused by disproportionally high transcription of a single 252 bp gene in the BGC. Following alignment against the nr database, these highly transcribed genes from the RiPP BGCs were determined to be the precursor peptides, with greatest similarity to nitrile hydratase like leader peptides (NHLPs), often referred to as proteusins (25, 26). We recovered a total of 343 precursor peptide gene sequences from all BGCs. Alignment and phylogenetic analysis of these sequences revealed 3 distinct clades (Fig. 9A). The largest clade of 191 sequences exhibited both the highest relative expression of precursor peptide genes (Fig. 9A) and very high levels of amino acid sequence conservation, particularly the last 34 residues at the C-terminal end of the sequence (Fig. 9B).

**Figure 9.**
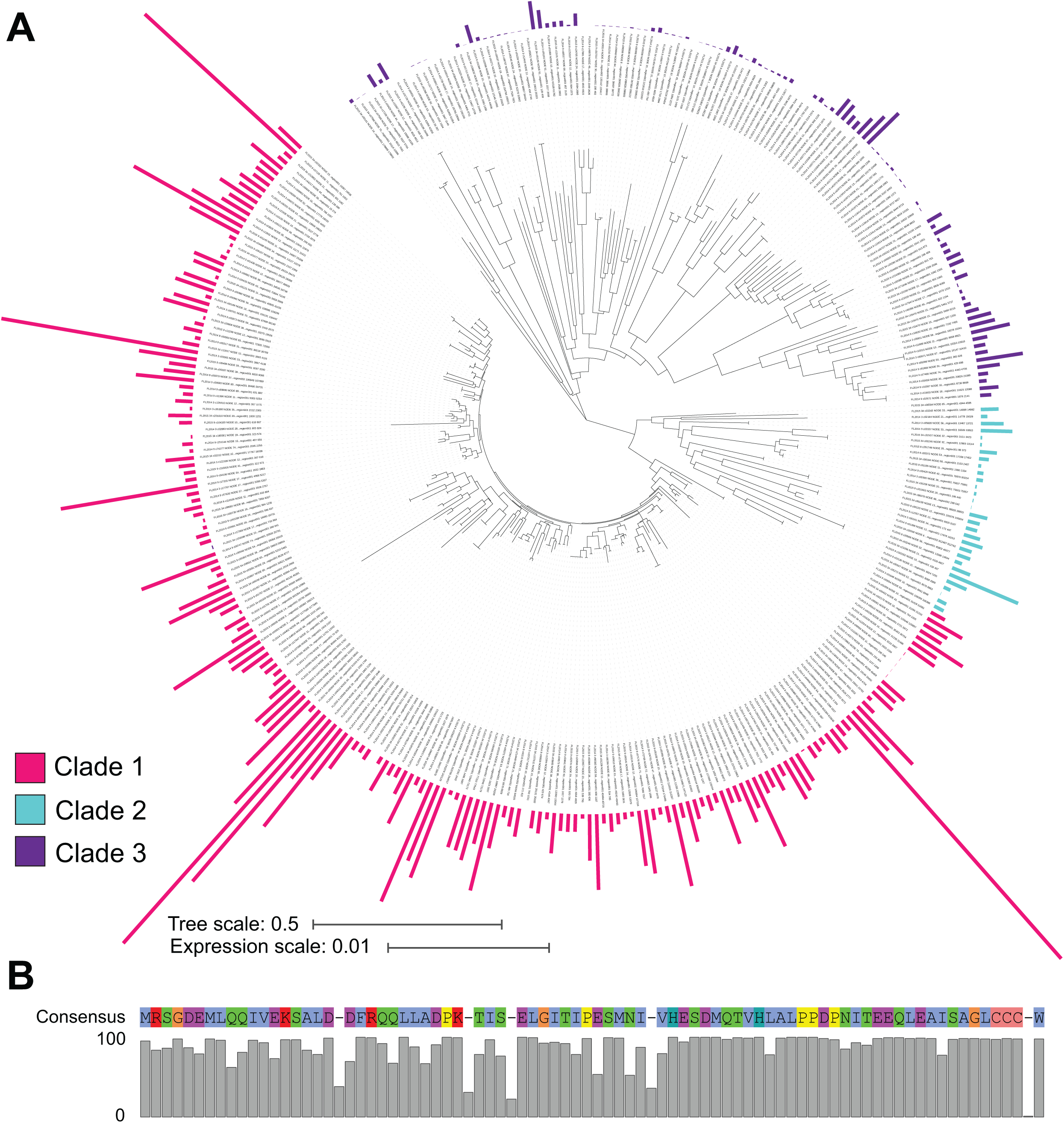
Expression of precursor peptide genes of 536 RiPP BGCs from four HMA sponges. **A.** Phylogenetic analysis of the protein sequence of the precursor peptide genes revealed three distinct clades wherein Clade 3 exhibited the highest expression levels. Expression levels of individual precursor peptide genes are indicated by bars and coloured according to clade. **B.** The precursor peptides from 191 BGCs across several different classes of sponge-associated bacteria exhibit a high level of sequence homology. Background colors of conserved residues follow Clustal X default coloring: Blue - Hydrophobic residues, Red - Positively charged residues, Magenta - Negatively charged residues, Green - Polar residues, Pink - Cysteines, Yellow - Prolines, Cyan - Aromatic residues.

These sequences appear to be most closely related to previously identified conserved Sponge Derived RiPP Proteusins (*srp*) genes. These *srp* genes are ubiquitous in various sponge species and are characterized by a LCCCW core peptide and the bromination of the terminal tryptophan residue, ultimately producing azol(in)e-containing RiPPs (26). Additionally, 116 of the precursor peptides found were co-located with genes encoding YcaO cyclodehydratases and 66 were co-located with genes encoding halogenases, lending further support that these gene clusters may produce similar compounds to those heterologously expressed by Ngyuen and colleagues (26). However, it appears that the YcaO recognition sequence identified by Ngyuyen and colleagues (LTEEQL) is different to the sequence identified in these bacteria, which appeared to have the consensus sequence of ITEEQL (Fig. 9B). Finally, Nguyen and colleagues determined that the BGCs are not localized to a single specialized symbiont but distributed throughout the sponge microbiome. The data presented here supports this and provides additional evidence that these BGCs are not just present but actively transcribed in a variety of bacterial phyla. A recent study by Loureiro and colleagues investigated the biosynthetic landscape of HMA sponges, and showed a high conservation of related RiPP BGCs indicating that these gene clusters may play an important role in the symbiosis between the host sponge and associated microbiome (27). Here we show that not only are these related RiPP BGCs widespread but they are being actively expressed, providing further evidence that these RiPP BGCs may produce compounds necessary for the function, survival or maintenance of the sponge-microbe holobiont.

## MATERIALS AND METHODS

### Sample Collection

Marine sponges were collected in July 2014 and August 2015 near the Keys Marine Lab (KML) in Key West, FL (n=34) from a depth of 2-5 m. Specifically, FL-20-3 and FL-20-9 were collected at LK point on July 14 2014 (N 24.80210 W 80.79451), FL2015-9 was collected behind KML on August 3 2015 (N 24.824549 W 80.816808), and FL2015-34 was collected at Conch Wall on August 4 2015 (N 24.79027 W 80.88528). Permission to collect these samples was granted through a Special Activity License issued by the Florida Fish and Wildlife Conservation Commission (SAL-15-1695-SR). A sample of sponge tissue (∼25 cm^3^) was placed in RNA*later*™ for metagenomic analysis within 10 min of collection, incubated at room temperature for 2.5 – 4.5 hours before long-term storage at -0 °C. The remaining tissue was frozen (-20 °C).

### Sample DNA/RNA extraction and sequencing

Both DNA and RNA were extracted from portions of sponge samples preserved in RNA*later*™ using previously described methods (28). In both cases the biological material was pulverized in a mortar and pestle under liquid nitrogen, followed either by DNA extraction through the use of proteinase K, SDS and CTAB followed by phenol-chloroform extraction, isopropanol precipitation, and re-purification by Genomic-tip 100/G (Qiagen) or RNA extraction with the use of the RNeasy Mini Kit (Qiagen). DNA was quantified by PicoGreen assay before shotgun Illumina HiSeq paired-end sequencing runs (2 × 101 bp and 2 × 126 bp). RNA was used to construct stranded Illumina libraries before paired-end sequencing runs (2 × 126 bp). 16S rRNA gene amplicon sequencing was carried out as previously described (28), using DNA extracted from both 2014 and 2015 sponge sample collections. All raw amplicon, metagenomic and metatranscriptomic data can be accessed under the BioProject accession PRJNA894551.

### Bacterial community analysis

16S rRNA gene sequence amplicon datasets for 21 *S. officinalis* and 13 *I. felix* sponge specimens were clustered into operational taxonomic units (OTUs) at a distance of 0.03 using mothur (29). Briefly, the paired-end read set for each sample was combined to create contigs. All sequences with ambiguous sequences, homopolymeric runs greater than 7, lengths smaller than 200 nucleotides and greater than 500 nucleotides were removed from the dataset. Chimeras were identified using VSEARCH (30) and removed from the datasets. The remaining sequences were then classified against the SILVA database (v.138) (31) and all sequences classified as “Chloroplast”, “Mitochondria”, ”unknown”, “Archaea” or “Eukaryota” were removed from the dataset. The remaining sequences were aligned to find conserved regions, and once again searched for chimeras and other contaminant sequences. Distances between sequences were calculated and sequences clustered into OTUs at a distance of 0.03 which is an approximation of bacterial species (32). A count table and associated fasta file were generated for these OTUs which were used for visualizations of bacterial community structure. Analysis of Similarity (ANOSIM) analysis was performed using the ANOSIM algorithm from the vegan (33) package in R. All commands and data used are available here: https://github.com/WiscEvan/sponge_paper (DOI: 10.6084/m9.figshare.23317025).

### Bacterial metagenomic analyses

Shotgun metagenomic reads were trimmed using Trimmomatic v.0.39 (34) and assembled using SPAdes v.3.14 (35). Resultant contigs were binned using Autometa (36) to generate metagenome-assembled genomes (MAGs). MAGs were manually curated using Automappa (https://github.com/WiscEvan/Automappa). MAG quality was assessed using CheckM v.1.1.3 (37) and taxonomically classified using GTDB-Tk v.1.4.1 (38) with database release 95 as reference. Gene prediction and annotation were performed using Prokka v.1.12 (39). Genes associated with primary metabolism were identified with KofamScan (40) with detailed output enabled. Only hits that were considered reliable (i.e. where the score is higher than the threshold value) were extracted from the results and considered for further analysis. Genes were similarly annotated against the COG database using eggNOG (41).

### Bacterial metatranscriptomic analysis

Transcripts were not trimmed as trimming has been shown to be both unnecessary and results in inaccurate expression estimates (42, 43). Instead, soft-clipping was enabled during alignment to metagenomic contigs. A pipeline (bacterial_metatranscriptomics.sh), which incorporates several custom Python scripts was implemented to achieve reproducible results in each sample and all scripts can be found at https://github.com/WiscEvan/sponge_paper/tree/master/sponge_paper/scripts.

Briefly, bacterial contigs were separated out from all assembled contigs using the pull_bacterial_contigs.py script and gene calling performed using Prodigal(44). Resultant general feature format files (*.gff) were converted to simplified tab-delimited text format files (*.gtf) using gffread (45). For each sponge sample, metagenomic reads and transcripts were separately aligned against the contigs using bbmap v. 38.18 (46) with softclipping enabled and sorted according to name using samtools v 1.9 (47). Counts of reads/transcripts per gene were enumerated using featureCounts (48), where read/transcript pairs were counted as a single mapped fragment, only paired reads/transcripts were counted, and reads were limited to mapping at a single location (i.e. multiple alignments were excluded). Validity of paired end distances were checked using default parameters. The raw counts were then normalized for gene size and sequencing depth using the reads per million (RPM) method. Briefly, the read or transcript counts per gene were divided by the length of the gene and multiplied by 1000 to calculate reads/transcripts per kilobase (RPK or TPK respectively). The RPK or TPK of all genes in all bacterial contigs (binned and unbinned) were then summed and divided by 1 million to produce the scaling factor. RPM values were calculated by dividing RPK by the scaling factor. RPM is an approximation for the abundance of a given gene. The same approach was used for the calculation of transcripts per million (TPM) from the metatranscriptomic data, which is an approximation of transcription levels of a given gene. Finally, the TPM was divided by the RPM to account for gene abundance i.e all expression levels are adjusted for genome bin abundance unless otherwise stated. Metadata per gene, such as inclusion into BGC clusters, KEGG annotations, and bin to which the gene belongs was then added to produce detailed tables and summaries. Please see the bacterial_metatranscriptomics.sh script and associated python scripts for detailed steps at https://github.com/WiscEvan/sponge_paper/tree/master/sponge_paper/scripts.

### Identification of biosynthetic gene clusters in all bacterial contigs

All contigs from all samples were searched for biosynthetic gene clusters using antiSMASH v 6.1.1. (49). Gene cluster families were identified using BiG-SCAPE (22) using –mix and –mibig options with mode set to auto and cutoffs set to 0.5.

### Host sponge mitogenomic analyses

Reference cytochrome C oxidase I subunit sequences were downloaded from NCBI for *Spongia officinalis* (YP_001648644.1)*, Ircinia felix* (JX306085.1, JX306086.1), *Neopetrosia proxima* (MK105443.1, MK105444.1, MK105445.1), *Spheciospongia vesparium* (MH749402.1) and *Tedania ignis* (AJ704976.1, AJ704977.1). Metagenome contigs were aligned to the corresponding downloaded reference mitogenomic fragments using the BLAST tblastx option (50). These tblastx searches were manually inspected for representative contig hits that fit a mitogenomic context. *Demospongia* mitogenomes have been observed to contain anywhere from two to twenty-seven tRNA and are relatively uniform in nucleotide content, with a range of thirteen to fifteen protein-coding genes (PCG) (51). Single putative mitogenome sequences were identified and were semi-automatically annotated using the Mitos2 web server (52) and verified using Artemis (53). Briefly, Mitos2 annotates each sequence using ARWEN (54), Mitfi (55), and tRNAscan-SE (56) aggregating results to construct a consensus about each tRNA and PCG region. These recovered and annotated mitogenomes were subjected to phylogenetic analysis for taxon validation of collected sponge samples.

### Host sponge taxon validation by mitogenome phylogeny

To assess placement of this study’s sponge mitogenomes within *Demospongia,* mitogenomes from *Plese et al.* (*57*) and *Lavrov et al.* (51) were downloaded. Protein coding genes were identified using MITOS2 and extracted into individually separated fasta files from each mitogenome. Nine protein-coding genes (ATP6, ATP8, ATP9, COX2, COX3, COB, NAD2, NAD3, NA5) were recovered from all genomes and bins and were separately concatenated, aligned and trimmed. The taxon-sorted and taxon-filtered gene sets were subjected to alignments using MAFFT v7.508 (58) with the -- auto option. Each of the gene alignments were trimmed using BMGE v1.12 (59) then combined into a nexus file using a custom script (concat_nex.py). A maximum-likelihood (ML) tree was constructed using IQ-Tree version 2.2.0.3 (60) with ModelFinder enabled (61). The final Bayesian inference tree was constructed using MrBayes 3.2.7a (62) with a starting tree specified using the IQ-Tree ML tree results. A burn-in percentage of 25%, with 4 chains (3 heated and one “cold”) was used. Due to initiating the analysis with the ML tree, chains were started using a slightly randomly perturbed version by setting nperts=4. A sampling frequency of 1000 was set with a rule to stop the run once the average standard deviation of split frequencies (ASDSF) dropped below 0.1. The maximum number of generations was set to 15,000,000. Trees were visualized on the interactive Tree of Life server (63) and rooted using *Rhizopus arrhizus* as an outgroup.

## ACKNOWLEDGEMENTS

SW was supported by the Gordon and Betty Moore Foundation (MMI-6920). CMC was supported by an NLM training grant to the Computation and Informatics in Biology and Medicine Training Program (NLM 5T15LM007359). We acknowledge support from NIAID (R21AI121704) as well as NSF (DBI 1845890). We thank the staff at the Keys Marine Laboratory for their assistance with the collection of the sponges. ChatGPT was used to suggest variations of the title of this manuscript to reduce verbosity.

## Figure and table legends

**Figure S1.** Average expression of COG annotated genes per bin where cumulative total expression is shown out of 100. Bins are organized by phyla, indicated by colored key, and faceted by sponge sample.

**Figure S2.** Complete network of all non-singleton BGC families in all four sponges sharing at least 50% homology. Each node represents a BGC. The shape of each node is indicative of the BGC type, the color is indicative of what phylum the host bin was classified as, and the size of the node is indicative of the average expression of the BGC.

**Figure S3.** All recovered BGCs that were classified as “Other” and expressed at relatively high levels. Recovered BGCs are on the left, while the network in which they were found is on the right. Predicted gene function is indicated with a colored key.

**Table S1.** Metadata associated with all recovered genome bins.

**Table S2.** Median gene expression of KEGG-annotated pathways in each sponge sample.

**Table S3.** Total percentage of genes annotated in each bin against the KEGG and COG databases.

**Table S4.** Median and average gene expression per COG category in each sponge sample.

**Table S5.** Total, average and median expression of each BGC identified, with associated metadata, in all four sponge samples.

**Table S6.** Metadata for individual genes in all BGCs classified within the RiPP BGC type.

## REFERENCES

1. Webster NS, Taylor MW. 2012. Marine sponges and their microbial symbionts: love and other relationships. Environ Microbiol 14:335–346.

2. Taylor MW, Radax R, Steger D, Wagner M. 2007. Sponge-associated microorganisms: evolution, ecology, and biotechnological potential. Microbiol Mol Biol Rev 71:295–347.

3. Moitinho-Silva L, Steinert G, Nielsen S, Hardoim CCP, Wu Y-C, McCormack GP, López-Legentil S, Marchant R, Webster N, Thomas T, Hentschel U. 2017. Predicting the HMA-LMA Status in Marine Sponges by Machine Learning. Front Microbiol 8:752.

4. Erwin PM, Coma R, López-Sendino P, Serrano E, Ribes M. 2015. Stable symbionts across the HMA-LMA dichotomy: low seasonal and interannual variation in sponge-associated bacteria from taxonomically diverse hosts. FEMS Microbiol Ecol 91:fiv115.

5. Sabrina Pankey M, Plachetzki DC, Macartney KJ, Gastaldi M, Slattery M, Gochfeld DJ, Lesser MP. 2022. Cophylogeny and convergence shape holobiont evolution in sponge–microbe symbioses. Nature Ecology & Evolution 6:750–762.

6. Kelly JB, Carlson DE, Low JS, Thacker RW. 2022. Novel trends of genome evolution in highly complex tropical sponge microbiomes. Microbiome 10:164.

7. Trindade-Silva AE, Rua C, Silva GGZ, Dutilh BE, Moreira APB, Edwards RA, Hajdu E, Lobo-Hajdu G, Vasconcelos AT, Berlinck RGS, Thompson FL. 2012. Taxonomic and functional microbial signatures of the endemic marine sponge Arenosclera brasiliensis. PLoS One 7:e39905.

8. Storey MA, Andreassend SK, Bracegirdle J, Brown A, Keyzers RA, Ackerley DF, Northcote PT, Owen JG. 2020. Metagenomic Exploration of the Marine Sponge Mycale hentscheli Uncovers Multiple Polyketide-Producing Bacterial Symbionts. mBio 11:e02997–19.

9. Yang Q, Cahn JKB, Piel J, Song Y-F, Zhang W, Lin H-W. 2022. Marine Sponge Endosymbionts: Structural and Functional Specificity of the Microbiome within Euryspongia arenaria Cells. Microbiol Spectr 10:e0229621.

10. Moitinho-Silva L, Díez-Vives C, Batani G, Esteves AIS, Jahn MT, Thomas T. 2017. Integrated metabolism in sponge–microbe symbiosis revealed by genome-centered metatranscriptomics. ISME J 11:1651–1666.

11. Campana S, Riesgo A, Jongepier E, Fuss J, Muyzer G, de Goeij JM. 2022. Meta-transcriptomic comparison of two sponge holobionts feeding on coral- and macroalgal-dissolved organic matter. BMC Genomics 23:674.

12. Morganti TM, Slaby BM, de Kluijver A, Busch K, Hentschel U, Middelburg JJ, Grotheer H, Mollenhauer G, Dannheim J, Rapp HT, Purser A, Boetius A. 2022. Giant sponge grounds of Central Arctic seamounts are associated with extinct seep life. Nat Commun 13:638.

13. Fiore CL, Labrie M, Jarett JK, Lesser MP. 2015. Transcriptional activity of the giant barrel sponge, Xestospongia muta Holobiont: molecular evidence for metabolic interchange. Front Microbiol 6:364.

14. Hudspith M, Rix L, Achlatis M, Bougoure J, Guagliardo P, Clode PL, Webster NS, Muyzer G, Pernice M, de Goeij JM. 2021. Subcellular view of host–microbiome nutrient exchange in sponges: insights into the ecological success of an early metazoan–microbe symbiosis. Microbiome 9:1–15.

15. Bayer K, Jahn MT, Slaby BM, Moitinho-Silva L, Hentschel U. 2018. Marine Sponges as Chloroflexi Hot Spots: Genomic Insights and High-Resolution Visualization of an Abundant and Diverse Symbiotic Clade. mSystems 3:e00150–18.

16. Kamke J, Sczyrba A, Ivanova N, Schwientek P, Rinke C, Mavromatis K, Woyke T, Hentschel U. 2013. Single-cell genomics reveals complex carbohydrate degradation patterns in poribacterial symbionts of marine sponges. ISME J 7:2287–2300.

17. Bowers RM, Kyrpides NC, Stepanauskas R, Harmon-Smith M, Doud D, Reddy TBK, Schulz F, Jarett J, Rivers AR, Eloe-Fadrosh EA, Tringe SG, Ivanova NN, Copeland A, Clum A, Becraft ED, Malmstrom RR, Birren B, Podar M, Bork P, Weinstock GM, Garrity GM, Dodsworth JA, Yooseph S, Sutton G, Glöckner FO, Gilbert JA, Nelson WC, Hallam SJ, Jungbluth SP, Ettema TJG, Tighe S, Konstantinidis KT, Liu W-T, Baker BJ, Rattei T, Eisen JA, Hedlund B, McMahon KD, Fierer N, Knight R, Finn R, Cochrane G, Karsch-Mizrachi I, Tyson GW, Rinke C, Genome Standards Consortium, Lapidus A, Meyer F, Yilmaz P, Parks DH, Eren AM, Schriml L, Banfield JF, Hugenholtz P, Woyke T. 2017. Minimum information about a single amplified genome (MISAG) and a metagenome-assembled genome (MIMAG) of bacteria and archaea. Nat Biotechnol 35:725–731.

18. Fieseler L, Quaiser A, Schleper C, Hentschel U. 2006. Analysis of the first genome fragment from the marine sponge-associated, novel candidate phylum Poribacteria by environmental genomics. Environ Microbiol 8:612– 624.

19. Fieseler L, Horn M, Wagner M, Hentschel U. 2004. Discovery of the novel candidate phylum “Poribacteria” in marine sponges. Appl Environ Microbiol 70:3724–3732.

20. Podell S, Blanton JM, Neu A, Agarwal V, Biggs JS, Moore BS, Allen EE. 2019. Pangenomic comparison of globally distributed Poribacteria associated with sponge hosts and marine particles. ISME J 13:468–481.

21. Guedes RLM, Prosdocimi F, Fernandes GR, Moura LK, Ribeiro HAL, Ortega JM. 2011. Amino acids biosynthesis and nitrogen assimilation pathways: a great genomic deletion during eukaryotes evolution. BMC Genomics 12 Suppl 4:S2.

22. Navarro-Muñoz JC, Selem-Mojica N, Mullowney MW, Kstring-namesar SA, Tryon JH, Parkinson EI, De Los Santos ELC, Yeong M, Cruz-Morales P, Abubucker S, Roeters A, Lokhorst W, Fernandez-Guerra A, Cappelini LTD, Goering AW, Thomson RJ, Metcalf WW, Kelleher NL, Barona-Gomez F, Medema MH. 2020. A computational framework to explore large-scale biosynthetic diversity. Nat Chem Biol 16:60–68.

23. Gilchrist CLM, Chooi Y-H. 2021. Clinker & clustermap.js: string-nameomatic generation of gene cluster comparison figures. Bioinformatics 37:2473–2475.

24. Gilchrist CLM, Booth TJ, van Wersch B, van Grieken L, Medema MH, Chooi Y-H. 2021. cblaster: A remote search tool for rapid identification and visualization of homologous gene clusters. Bioinformatics Advances 1:vbab016.

25. Haft DH, Basu MK, Mitchell DA. 2010. Expansion of ribosomally produced natural products: a nitrile hydratase- and Nif11-related precursor family. BMC Biology 8:70.

26. Nguyen NA, Lin Z, Mohanty I, Garg N, Schmidt EW, Agarwal V. 2021. An Obligate Peptidyl Brominase Underlies the Discovery of Highly Distributed Biosynthetic Gene Clusters in Marine Sponge Microbiomes. J Am Chem Soc 143:10221–10231.

27. Loureiro C, Galani A, Gavriilidou A, Chaib de Mares M, van der Oost J, Medema MH, Sipkema D. 2022. Comparative Metagenomic Analysis of Biosynthetic Diversity across Sponge Microbiomes Highlights Metabolic Novelty, Conservation, and Diversification. mSystems 7:e0035722.

28. Miller IJ, Weyna TR, Fong SS, Lim-Fong GE, Kwan JC. 2016. Single sample resolution of rare microbial dark matter in a marine invertebrate metagenome. Sci Rep 6:34362.

29. Schloss PD, Westcott SL, Ryabin T, Hall JR, Hartmann M, Hollister EB, Lesniewski RA, Oakley BB, Parks DH, Robinson CJ, Sahl JW, Stres B, Thallinger GG, Van Horn DJ, Weber CF. 2009. Introducing mothur: Open- source, platform-independent, community-supported software for describing and comparing microbial communities. Appl Environ Microbiol 75:7537–7541.

30. Rognes T, Flouri T, Nichols B, Quince C, Mahé F. 2016. VSEARCH: A versatile open source tool for metagenomics. PeerJ 4:e2584.

31. Quast C, Pruesse E, Yilmaz P, Gerken J, Schweer T, Yarza P, Peplies J, Glöckner FO. 2013. The SILVA ribosomal RNA gene database project: improved data processing and web-based tools. Nucleic Acids Res 41:D590–6.

32. Chappidi S, Villa EC, Cantarel BL. 2019. Using Mothur to Determine Bacterial Community Composition and Structure in 16S Ribosomal RNA Datasets. Curr Protoc Bioinformatics 67:e83.

33. Dixon P. 2003. VEGAN, a package of R functions for community ecology. J Veg Sci 14:927–930.

34. Bolger AM, Lohse M, Usadel B. 2014. Trimmomatic: a flexible trimmer for Illumina sequence data. Bioinformatics 30:2114–2120.

35. Bankevich A, Nurk S, Antipov D, Gurevich AA, Dvorkin M, Kulikov AS, Lesin VM, Nikolenko SI, Pham S, Prjibelski AD, Pyshkin AV, Sirotkin AV, Vyahhi N, Tesler G, Alekseyev MA, Pevzner PA. 2012. SPAdes: A new genome assembly algorithm and its applications to single-cell sequencing. J Comput Biol 19:455–477.

36. Miller IJ, Rees ER, Ross J, Miller I, Baxa J, Lopera J, Kerby RL, Rey FE, Kwan JC. 2019. string-nameometa: string-nameomated extraction of microbial genomes from individual shotgun metagenomes. Nucleic Acids Res 47:e57.

37. Parks DH, Imelfort M, Skennerton CT, Hugenholtz P, Tyson GW. 2015. CheckM: assessing the quality of microbial genomes recovered from isolates, single cells, and metagenomes. Genome Res 25:1043–1055.

38. Chaumeil P-A, Mussig AJ, Hugenholtz P, Parks DH. 2020. GTDB-Tk: a toolkit to classify genomes with the Genome Taxonomy Database. Bioinformatics 36:1925–1927.

39. Seemann T. 2014. Prokka: Rapid prokaryotic genome annotation. Bioinformatics 30:2068–2069.

40. Aramaki T, Blanc-Mathieu R, Endo H, Ohkubo K, Kanehisa M, Goto S, Ogata H. 2020. KofamKOALA: KEGG ortholog assignment based on profile HMM and adaptive score threshold. Bioinformatics 36:2251–2252.

41. Huerta-Cepas J, Szklarczyk D, Heller D, Hernández-Plaza A, Forslund SK, Cook H, Mende DR, Letunic I, Rattei T, Jensen LJ, von Mering C, Bork P. 2019. eggNOG 5.0: a hierarchical, functionally and phylogenetically annotated orthology resource based on 5090 organisms and 2502 viruses. Nucleic Acids Res 47:D309–D314.

42. Williams CR, Baccarella A, Parrish JZ, Kim CC. 2016. Trimming of sequence reads alters RNA-Seq gene expression estimates. BMC Bioinformatics 17:103.

43. Liao Y, Shi W. 2020. Read trimming is not required for mapping and quantification of RNA-seq reads at the gene level. NAR Genom Bioinform 2:lqaa068.

44. Hyatt D, Chen G-L, Locascio PF, Land ML, Larimer FW, Hauser LJ. 2010. Prodigal: prokaryotic gene recognition and translation initiation site identification. BMC Bioinformatics 11:119.

45. Pertea G, Pertea M. 2020. GFF Utilities: GffRead and GffCompare. F1000Research https://doi.org/10.12688/f1000research.23297.2.

46. Bushnell B. 2014. BBMap: a fast, accurate, splice-aware aligner. Lawrence Berkeley National Lab.(LBNL), Berkeley, CA (United States).

47. Li H, Handsaker B, Wysoker A, Fennell T, Ruan J, Homer N, Marth G, Abecasis G, Durbin R, 1000 Genome Project Data Processing Subgroup. 2009. The Sequence Alignment/Map format and SAMtools. Bioinformatics 25:2078–2079.

48. Liao Y, Smyth GK, Shi W. 2014. featureCounts: an efficient general purpose program for assigning sequence reads to genomic features. Bioinformatics 30:923–930.

49. Blin K, Shaw S, Kloosterman AM, Charlop-Powers Z, van Wezel GP, Medema MH, Weber T. 2021. antiSMASH 6.0: improving cluster detection and comparison capabilities. Nucleic Acids Res 49:W29–W35.

50. Altschul SF, Gish W, Miller W, Myers EW, Lipman DJ. 1990. Basic local alignment search tool. J Mol Biol 215:403–410.

51. Wang X, Lavrov DV. 2008. Seventeen new complete mtDNA sequences reveal extensive mitochondrial genome evolution within the Demospongiae. PLoS One 3:e2723.

52. Donath A, Jühling F, Al-Arab M, Bernhart SH, Reinhardt F, Stadler PF, Middendorf M, Bernt M. 2019. Improved annotation of protein-coding genes boundaries in metazoan mitochondrial genomes. Nucleic Acids Res 47:10543–10552.

53. Rutherford K, Parkhill J, Crook J, Horsnell T, Rice P, Rajandream MA, Barrell B. 2000. Artemis: sequence visualization and annotation. Bioinformatics 16:944–945.

54. Laslett D, Canbäck B. 2008. ARWEN: a program to detect tRNA genes in metazoan mitochondrial nucleotide sequences. Bioinformatics 24:172–175.

55. Jühling F, Pütz J, Bernt M, Donath A, Middendorf M, Florentz C, Stadler PF. 2012. Improved systematic tRNA gene annotation allows new insights into the evolution of mitochondrial tRNA structures and into the mechanisms of mitochondrial genome rearrangements. Nucleic Acids Res 40:2833–2845.

56. Lowe TM, Eddy SR. 1997. tRNAscan-SE: a program for improved detection of transfer RNA genes in genomic sequence. Nucleic Acids Res 25:955–964.

57. Plese B, Kenny NJ, Rossi ME, Cárdenas P, Schuster A, Taboada S, Koutsouveli V, Riesgo A. 2021. Mitochondrial evolution in the Demospongiae (Porifera): Phylogeny, divergence time, and genome biology. Mol Phylogenet Evol 155:107011.

58. Katoh K, Standley DM. 2013. MAFFT multiple sequence alignment software version 7: improvements in performance and usability. Mol Biol Evol 30:772–780.

59. Criscuolo A, Gribaldo S. 2010. BMGE (Block Mapping and Gathering with Entropy): a new software for selection of phylogenetic informative regions from multiple sequence alignments. BMC Evol Biol 10:210.

60. Nguyen L-T, Schmidt HA, von Haeseler A, Minh BQ. 2015. IQ-TREE: a fast and effective stochastic algorithm for estimating maximum-likelihood phylogenies. Mol Biol Evol 32:268–274.

61. Kalyaanamoorthy S, Minh BQ, Wong TKF, von Haeseler A, Jermiin LS. 2017. ModelFinder: fast model selection for accurate phylogenetic estimates. Nat Methods 14:587–589.

62. Ronquist F, Huelsenbeck JP. 2003. MrBayes 3: Bayesian phylogenetic inference under mixed models. Bioinformatics 19:1572–1574.

63. Letunic I, Bork P. 2021. Interactive Tree Of Life (iTOL) v5: an online tool for phylogenetic tree display and annotation. Nucleic Acids Res 49:W293–W296.

